# Regulation of ADP-ribosyltransferase activity by ART domain dimerization in PARP15

**DOI:** 10.1101/2024.04.04.588081

**Authors:** Carmen Ebenwaldner, Antonio Ginés García Saura, Simon Ekström, Katja Bernfur, Herwig Schüler

## Abstract

PARP15 is a mono-ADP-ribosyltransferase with unknown functions. Its evolutionary relationship with PARP14 suggests roles in antiviral defense; its ability to modify RNA and localization to stress granules point to functions in the regulation of translation. PARP15 also modifies itself and other proteins using its ADP-ribosyltransferase (ART) domain and contains two macrodomains predicted to bind ADP-ribosyl on targets. We used biochemical and biophysical analysis to study how the ADP-ribosyltransferase activity of PARP15 is regulated. Here we show that the catalytic domain of PARP15 dimerizes with mid-nanomolar affinity, forming the same dimer interface in solution that had already been captured by X-ray crystallography of the domain. Furthermore, we show that the formation of dimers is a prerequisite for catalytic activity and that monomeric mutant variants of the domain were catalytically inactive. Our findings suggest a regulatory mechanism by which dimerization is linked to either target engagement or placement of a catalytic residue, rather than NAD+ co-substrate binding, and by which the two protomers of the dimer operate independent of one another. Together, our results uncover a novel mechanism of regulation in a PARP family enzyme, which might inspire new avenues of pharmacological intervention.

## Introduction

Proteins of the poly(ADP-ribosyl) polymerase – short PARP – family are defined by a common, structurally highly conserved catalytic ADP-ribosyltransferase (ART) domain [1]. ADP-ribosylation, in which the ADP-ribose moiety from NAD+ is transferred onto target molecules, partakes in the regulation of processes including DNA damage repair, transcription and translation, protein homeostasis, and antiviral defense [2-4]. ADP-ribosylation is of prime therapeutic interest due to its contribution to DNA damage repair and de-regulated signaling pathways in cancers [5,6]. Mechanisms that regulate PARP enzymatic activities are bound to be different for every PARP family member, as these differ in the types and numbers of accessory domains. Such domains include recognition modules for DNA, RNA, ADP-ribosylated macromolecules, and protein interaction modules [4]. Currently, the functioning of PARP accessory domains in concert with the ART domains is enigmatic with one exception: We now have a rather clear picture of how a DNA strand break binding event is relayed, via internal domains, to the ART domain to boost its activity [7-10].

The subfamily of macrodomain containing PARP enzymes (PARP9, PARP14 and PARP15) was originally identified as B-aggressive lymphoma (BAL) proteins 1-3 [11]. PARP9 and PARP14 contain a macrodomain with ADP-ribosyl glycohydrolase activity [12-14] as well as one and two ADP-ribosyl binding macrodomains, respectively [15-17]. PARP15, the third member of the subfamily, originated by partial duplication of the structurally more complex PARP14 early during mammalian evolution. Both genes have remained under positive selection pressure; but uniquely, the parp15 gene was lost again independently in some lineages, and truncated or duplicated in others [18]. Thus, evolutionary analysis strongly ties PARP15 to virus defence [18]. Humans possess two isoforms of PARP15; isoform-1 contains an unstructured N-terminal extension of unknown function, two macrodomains, and a C-terminal ART domain, whereas isoform-2 features a shorter N-terminal extension and lacks macrodomain-1 but is otherwise identical to isoform-1 [18]. Macrodomain-1 is poorly characterized, whereas crystal structures and biochemical analyses suggest that macrodomain-2 is an ADP-ribosyl binding module [15,16]. PARP15 has been shown to localize to stress granules [19], cytoplasmic condensates consisting of proteins and RNA that form upon exposure to a variety of stressors, including viral infection [20]. More recently, PARP15 was found to ADP-ribosylate 5’phosphorylated RNA ends *in vitro* and in cells [21,22].

Despite the recent evolutionary kinship of PARP14 and PARP15, the human PARP15 ART domain has higher affinity for NAD+ and modifies more sites on itself *in vitro* [23,24]. Here, we set out to study the molecular basis of these differences. We show that the PARP15 ART domain forms a dimer in solution and that dimer formation is required for MARylation activity. We show that the dimer interface in solution is identical with the interface that has been observed by X-ray crystallography. Mutation of a central salt bridge network formed by Arg576 and Asp665 abrogated dimerization, yielding a catalytically inactive monomer. Finally, we find that the activation by dimerization of one protomer is not dependent on a functional catalytic site in the second protomer. In summary, this study proposes catalytic domain homodimerization as a novel regulatory mechanism for a member of the PARP enzyme family.

## Results

During routine purification of the PARP15 ART domain (Asn^481^-Ala^678^), we observed that its elution profile in size exclusion chromatography was inconsistent with a monomer of the expected size. To characterize this behavior, we carried out size exclusion chromatography in combination with right-angle and low-angle light scattering measurements. The results indicated that the PARP15 ART domain eluted as a dimer (Fig. 1A). For comparison, we characterized several protein domains related to PARP15. The PARP14 ART domain (His^1608^-Lys^1801^; 65% sequence identity) was monomeric and the PARP10 ART domain (N^819^-V^1007^; 37% sequence identity) was largely monomeric (Fig. 1B,C). A PARP15 macrodomain-2-ART domain construct (m2-ART; Gly^287^-Ala^678^) eluted in a broad peak containing both monomeric and dimeric species (Fig. 1D) although macrodomain-2 alone (m2; Gly^287^Asn^470^) eluted as a monomer (Fig. 1E). To substantiate the interpretation that the PARP15 ART domain was a dimer in solution, we treated proteins with BS^3^, a 11.4 Å chemical crosslinker [25]. Using a half-molar equivalent concentration of crosslinker per protein, PARP15 ART and m2-ART could be crosslinked as dimers, whereas PARP15 m2 and PARP14 ART could not (Fig. 1G). This supported the previous observation that the PARP15 ART domain was a dimer in solution.

**Figure 1.**
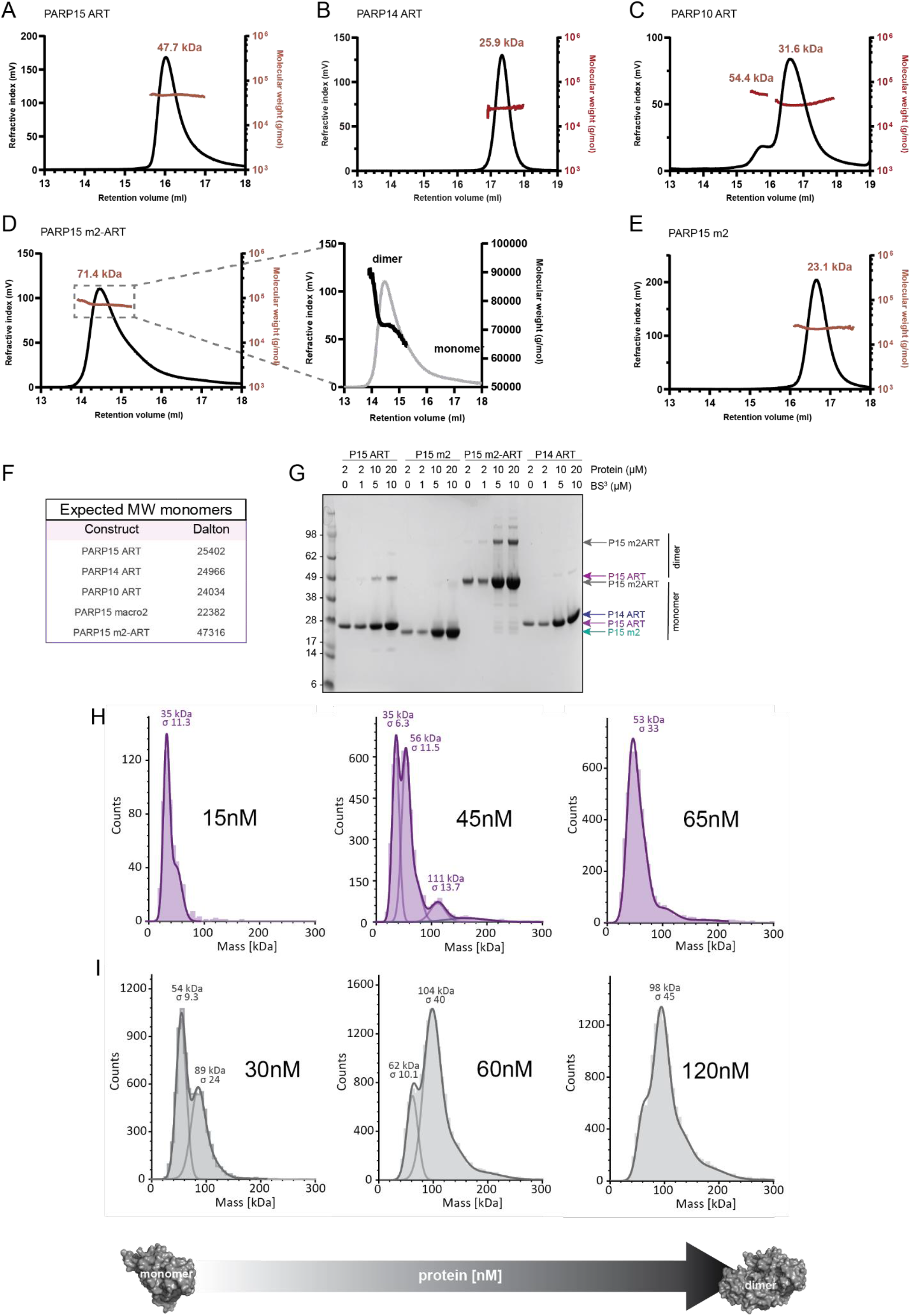
PARP15 dimerizes via its ART domain. **(A-F)** SEC-RALS/LALS analysis reveals that the PARP15 ART domain elutes as a dimer (A) on the size exclusion column whereas the ART domains of PARP14 (B) and PARP10 (C) are monomeric. The peak of the PARP15 m2-ART construct (D) contains both monomeric and dimeric species. The PARP15 macrodomain-2 (E) is a monomer. Further details are given in Supplementary Table S1. (F) Expected molecular weights of the respective monomers. **(G)** PARP15 ART and m2-ART constructs show dimers on an SDS-PAGE gel after crosslinking with BS^3^. PARP15 m2 and PARP14 ART do not form dimers. **(H**,**I)** Size distribution of single particles of PARP15 ART (H; purple) and PARP15 m2-ART (I; grey) in the nanomolar concentrations range, observed by mass photometry.

To gain more insight into the physical properties of the PARP15 ART domain dimer, we performed single particle sizing experiments using mass photometry. This analysis suggested that ART and m2-ART constructs were monomeric at low nanomolar concentrations, and the size distributions shifted toward dimers with increasing protein concentrations in a range up to 120 nM (Fig. 1H,I). To establish the affinity of the ART domain dimer interaction, we employed fluorescence polarization. Fluorescein-labelled PARP15 ART domain was kept constant at either 1 or 10 nM, and fluorescence anisotropy was monitored over increasing concentrations of unlabelled ART domain. The resulting data were fitted to a one-site binding term, giving dissociation constants of 258 and 273 nM, respectively (Fig. 2).

**Figure 2.**
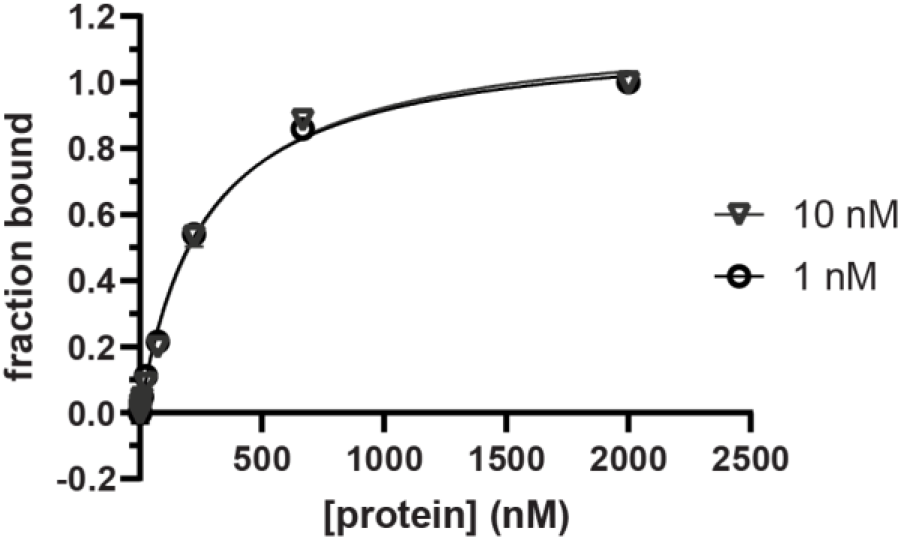
PARP15 dimerizes with a dissociation constant in the nanomolar range. Unlabelled ART domain was titrated against fluorescein-labelled ART domain at two concentrations, and the fraction of ART domain dimer was determined by fluorescence polarization spectroscopy. The data suggest a K_d_ for the dimer interaction of 258 and 273 nM, respectively.

Having established that the PARP15 ART domain is a dimer in solution, we were interested in defining the location of the dimer interface. Abundant crystallographic data is available for this domain in the Protein Data Bank and with no exception, the domain crystallized featuring a homodimer in the asymmetric unit [26-28]. An analysis of 39 crystal structures using the PISA server [29] gave a mean buried surface area of 1046±23 Å^2^ for the crystallographic dimer interface. For comparison, a crystal structure of the homodimeric 14-3-3β (PDB id 2BQ0) [30] features a common buried surface area between the protomers of 1030 Å^2^. Thus, it was reasonable to assume that the dimer interface present in the PARP15 ART domain crystals may be relevant to the dimer in solution. In all PARP15 ART domain models based on crystallographic data, the dimer interface was made up of side chains in helix α2 of each monomer, as well as a prominent salt bridge network formed by residues in the two turns following helix α3 and between strands β10 and β11, respectively from each monomer (Fig. 3A-C and Supplementary Table S2). In addition, hydrogen bonds and hydrophobic interactions were formed across the interface by side chains in the vicinities of the sites above. AlphaFold2-multimer [31] predicted a dimer with high confidence (pTM and ipTM scores of > 0.9), with an interface identical to the one in the crystallographic dimers (Fig. 3A and Supplementary Figure S1).

**Figure 3.**
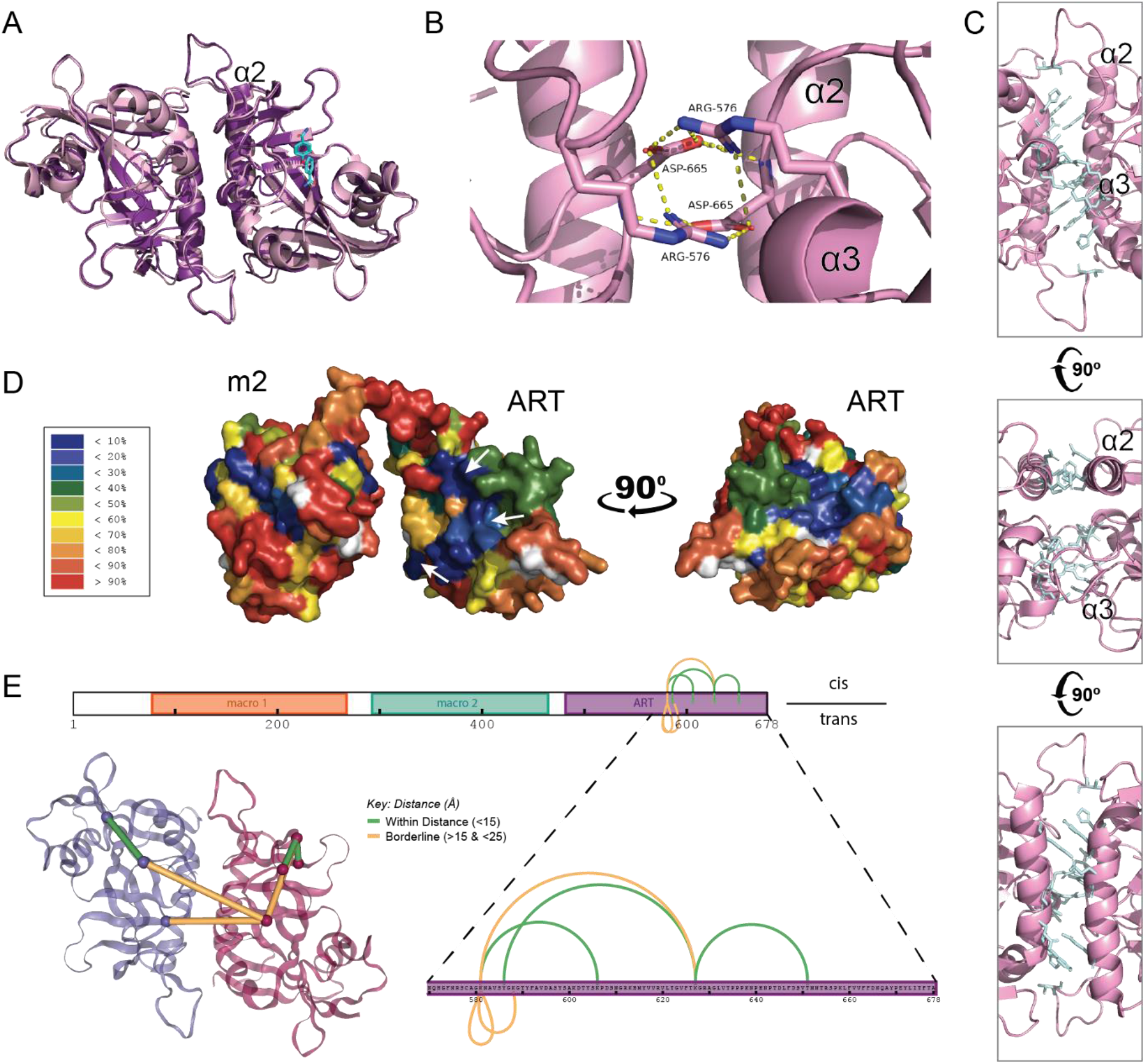
A combination of experimental and computational tools defines the dimer interface. (**A-C**) A structural model generated by AlphaFold2 multimer suggests an ART domain dimer (dark violet) that superimposes with an RMSD(C_α_) of 0.876 Å with the crystal dimer (6ry4 [28]; light pink). The ligand (cyan) is bound to the active site of one protomer in the crystal structure. (**B**) The dimer interface contains a prominent salt bridge network formed by side chains Arg^576^ and Asp^665^ of each protomer. (**C**) Helixes α2 of each protomer as well as residues in the vicinity of α3 interact at the dimer interface. (**D**) AlphaFold model of PARP15 m2-ART, colored for protection against hydrogen exchange at 2.5 h incubation in D_2_O (red, all hydrogens exchanged; blue, all hydrogens protected). Detailed illustrations of the HDX-MS data are shown in Supplementary Fig. S2. (**E**) BS^3^-crosslinked ART domain peptides identified by mass spectrometry are in agreement with the dimer interface observed by X-ray crystallography. Crosslinked peptide sequences and masses are given in Supplementary Table S3.

To test whether the crystallographic dimer interface was present in the dimer in solution, we analyzed the m2-ART construct using hydrogen-deuterium exchange coupled with mass spectrometry (HDX-MS) [32]. The results showed that a surface that coincides with the crystallographic dimer interface is fully protected from deuterium exchange during the experiment (2.5 hours; Fig. 3D and Supplementary Fig. S2). A second patch on the ART domain and a ridge over the surface of macrodomain-2 were also protected indicating that macrodomain-2 might contribute to the dimer interaction. Furthermore, we crosslinked the PARP15 ART domain under the conditions of Fig. 1G and analyzed the product by mass spectrometry after proteolytic cleavage. This analysis revealed six high-confidence putative crosslinks, two of which unambiguously represent inter-protomer crosslinks as they were identified on overlapping peptides (Supplementary Table S3). When mapped on the structure of the crystallographic dimer, all six crosslinks are in agreement with the distance constraints of the BS^3^crosslinker (Fig. 3E). From these results we conclude that the crystallographic dimer interface is highly similar to or identical with the dimer interface in solution.

Protein oligomerization is a common means of regulation. Therefore, it was important to investigate whether PARP15 catalytic activity was related to its oligomerization state. To that end, we created several ART domain variants in which either of the two residues R576 and D665, which participate in the interface salt bridge network (Fig. 3B), had been mutated (Fig. 4A). We made variants with three different substitutions for R576 and two substitutions for D665. Each mutant protein construct showed a higher retention volume than the wild type construct in analytical size exclusion chromatography (Fig. 4B), indicating that they were monomeric. All dimer interface mutants eluted in one symmetric peak except for R576A, which may retain partial dimerization ability (Fig. 4B). Then, we incubated the dimer interface mutant proteins, alone or in pairwise combinations, with NAD+ (10% biotin-NAD+) and analyzed the reactions by Western blotting, followed by detection of auto-MARylation levels using HRP-streptavidin (Fig. 4C). Alternatively, we analyzed catalytic reactions containing no biotin-NAD+ using a GFP-macrodomain overlay assay [33]. All mutant variants were less active than the wild type construct (Fig. 4C,D). The R576A mutant retained the highest level of activity, consistent with the observation of residual dimerization (Fig. 4B). The charge reversion mutants R576D and R576E had no measurable catalytic activity, although circular dichroism spectroscopy indicated they were properly folded (Supplementary Figure S3). The combination of equal amounts of the charge reversion mutants at R576 and at D665 did not result in rescue of MARylation activity. The combination of equal amounts of the charge-to alanine mutants at R576 and at D665, however, retained low levels of activity (Fig. 4D), again likely due to weak or transient dimer formation as in the R576A mutant alone.

**Figure 4.**
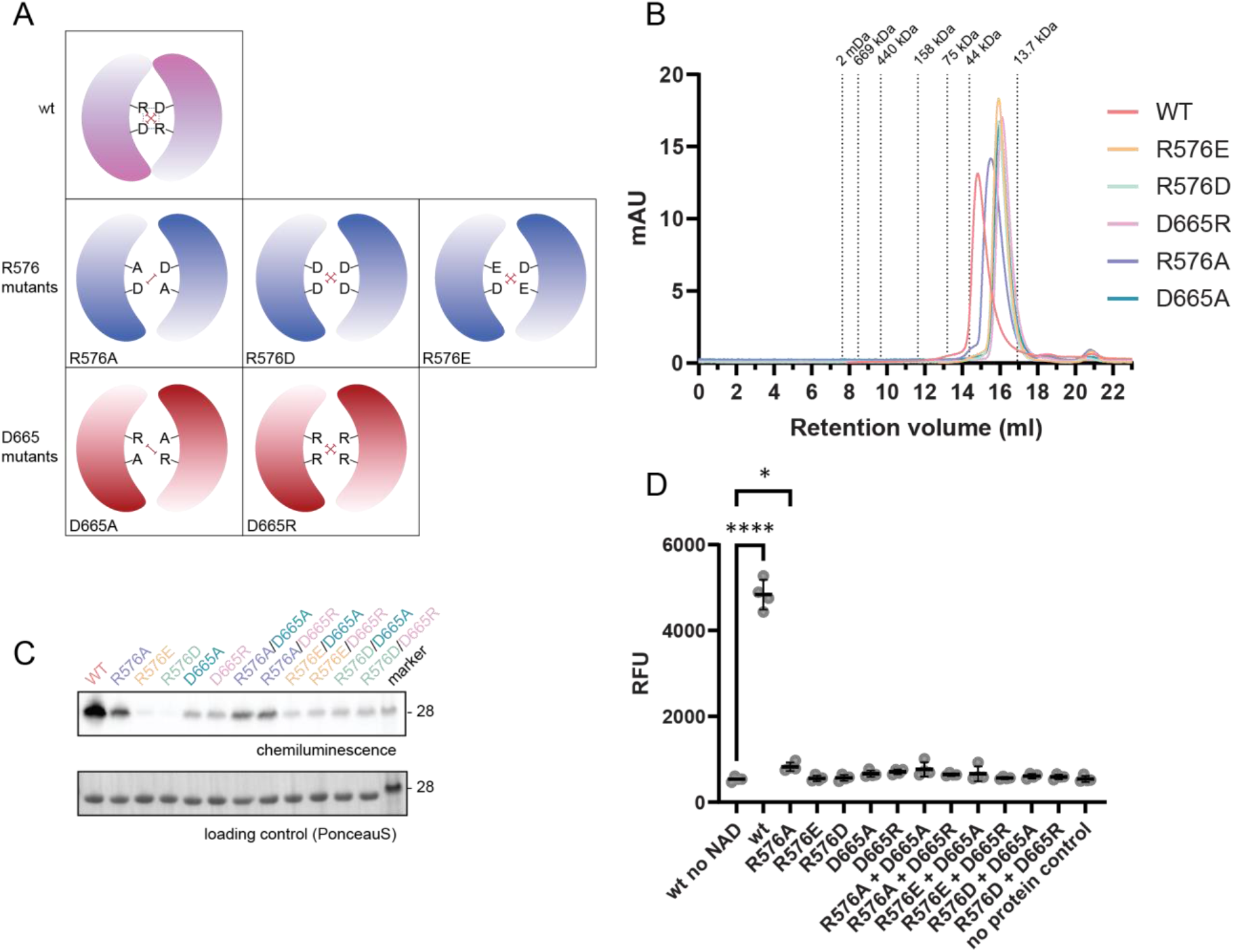
Point mutations at the dimer interface disrupt dimerization and catalytic activity. (**A**) Schematic representation of the wild type dimer with its central salt bridge network consisting of one arginine and one aspartate per protomer. The point mutations introduced are illustrated below, with the expected electrostatic repulsions indicated as red dashes. (**B**) All interface mutant variants eluted as monomers in analytical size exclusion chromatography. (**C**,**D**) All mutant constructs were significantly less active compared to wt PARP15, as shown by Western blot analysis of biotin-NAD+ spiked reactions (C; 10 µM total enzyme) and by a fluorescent macrodomain overlay assay (D; 1 µM total enzyme). The combination of variants with different surface mutations could not rescue the loss-of-activity phenotype.

To further explore the link between PARP15 catalytic activity and dimer formation, we measured auto-modification activity over a range of enzyme concentrations below and above the dissociation constant for PARP15 dimer formation, for the ART domains of the monomeric PARP10 (Asn^819^-Val^1007^) and of PARP15. Fig. 4A shows that PARP10 ART domain auto-modification increased linearly over the entire concentration range. In contrast, PARP15 ART domain activity displayed a sharp incline as the enzyme concentration approached the dissociation constant for dimerization. These results confirm that dimer formation is a prerequisite for PARP15 MARylation activity. To strengthen this conclusion, we repeated the experiment with the PARP15 dimer interface mutant D665R (Fig. 5A). This monomeric variant remained inactive over the range of concentrations tested.

**Figure 5.**
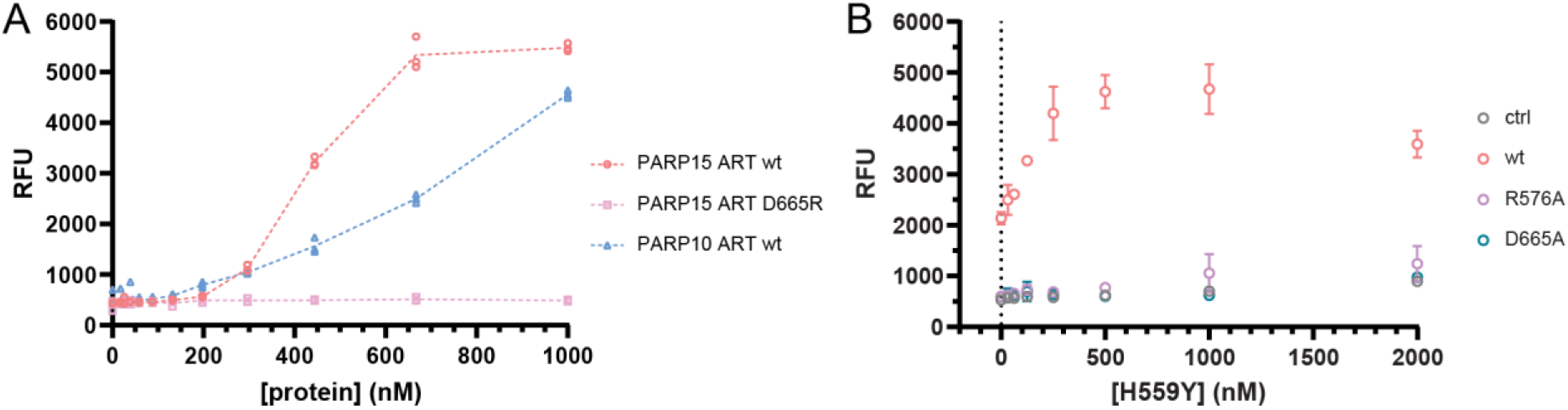
One active protomer is sufficient for catalytic activity in the PARP15 dimer. (**A**) Extent of auto-modification in PARP15 and PARP10 ART domain constructs over a range of enzyme concentrations. Whereas PARP10 activity showed a near-linear increase of activity with increasing protein concentration, PARP15 activity increase sharply at a concentration near the dissociation constant of dimer formation. (**B**) PARP15 macrodomain-2 was bound to high-binding 96-well plates to serve as target for ADP-ribosylation. Wild type PARP15 ART, D665A mutant, or R576A mutant, were kept at a concentration around the dissociation constant (250 nM) of wt PARP15 and increasing concentrations of the catalytically inactive mutant H559Y were added. Wild type and mutant ART were washed out and ADP-ribosylation on macrodomain-2 was quantified, revealing that the addition of catalytically dead mutant boosts the activity of the wt PARP15, but does not rescue the activity of the two dimer mutants. In the control, only H559Y mutant is present to show that it does not have any activity over m2 by itself.

Having established that the catalytically active species of the PARP15 ART domain is a dimer, we asked whether each protomer within the dimer must be catalytically active for the reactions to proceed. For that purpose, we coated assay plates with PARP15 macrodomain-2 as target for MARylation. Either wild type PARP15 ART domain or the monomeric variants D665A and R576A were added at a concentration of 250 nM, close to the dissociation constant of the wild type dimer. Finally, the total concentration of ART domains was titrated with the catalytically inactive PARP15 ART domain mutant H559Y. Wild type ART domain alone at 250 nM caused a moderate level of MARylation (Fig. 5B). Addition of inactive H559Y mutant caused a rise in MARylation levels, which reached saturation around 500 nM total enzyme (sum of wild type and H559Y mutant; Fig. 5B). When either of the dimerization mutants R576A or D665A were present and H559Y mutant was added, no rise in MARylation activity was observed. We conclude that the H559Y mutant construct can participate in heterodimer formation with the wild type ART domain, thus activating catalysis without processing NAD+. This implies that catalytic activity in each protomer within the dimer is independent of the activity in the second protomer.

This experiment (Fig. 5B) showed that only one protomer within the dimer needs to be catalytically competent in order for the dimer to display MARylation activity. We also addressed the question whether dimerization was needed for ligand binding in the active site. First, we used differential scanning fluorimetry to assess the thermal stability of the ART domain. All dimer interface mutations except D665A lowered the T_m_ of the ART domain (Fig. 6A) and all proteins melted showing symmetric transitions (Supplementary Fig. S4). Then, we used the same assay to probe the ability of the ART domain to bind 3-aminobenzamide (an analog of nicotinamide, the product of the MARylation reaction). All dimer interface mutant constructs were stabilized by 3-aminobenzamide to roughly the same degree as the wild type construct (Fig. 6A,B). This was the case also for R576E and R576D, which showed no indication of either dimer formation or MARylation activity. The active-site mutant H559Y, that is ligand binding incompetent [26], was only slightly stabilized by 3-aminobenzamide. We conclude that the active site of the PARP15 ART domain is competent for ligand binding in the active site independent of its oligomerization state.

**Figure 6.**
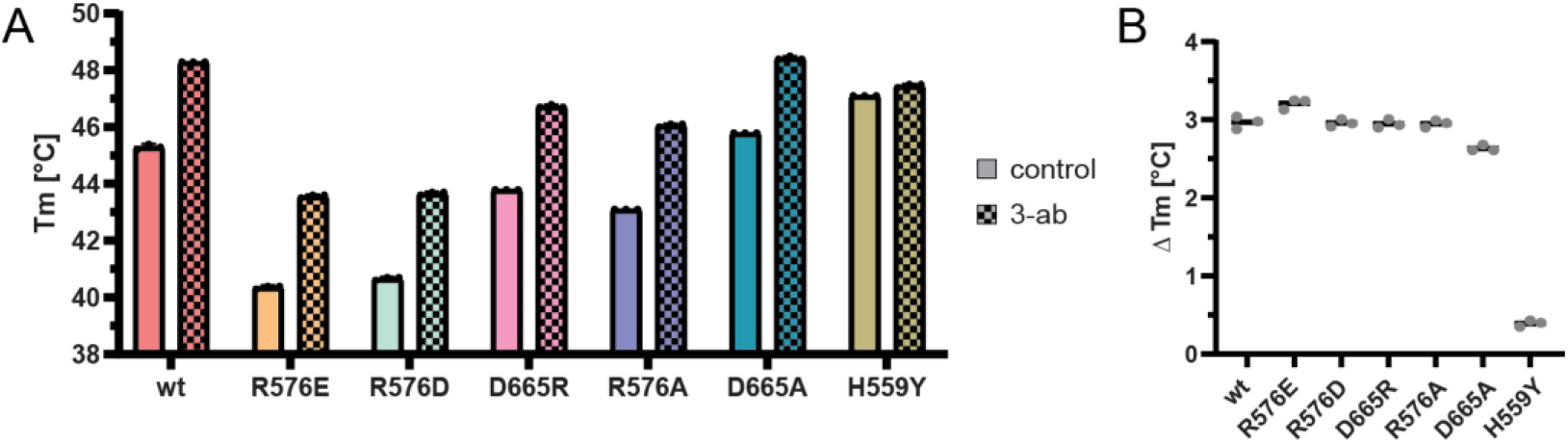
Point mutations at the dimer interface do not disrupt ligand binding at the active site. (**A**) Representation of the melting temperature (T_m_) of the PARP15 ART domain, and the effect of surface mutations and excess 3-aminobenzamide. (**B**) Shifts upon 3-aminobenzamide binding in the melting temperatures (ΔT_m_) of wild type and mutant ART domain.

## Discussion

The ART domain is the defining feature of the PARP family. Its overall fold is highly conserved, with well-defined key amino acids that position the NAD+ molecule in a strained orientation, enabling its activation, hydrolysis, and attachment of the ADP-ribose moiety onto an acceptor molecule [34]. There is currently no evidence for the need of any accessory domain for the ART domain to be catalytically active. However, PARP1 and PARP2 have an α-helical subdomain in direct contact with their ART domain. In PARP1, the ART domain is kept in a catalytically inactive state unless a DNA binding event is relayed throughout the multidomain assembly to the α-helical subdomain, which undergoes a conformational change to activate the ART domain [7,8,10]. A similar mechanism regulates PARP2 [9]. Homologous helical extensions to the ART domain in PARP3 [35] and PARP4 [36] and a non-homologous α-helical subdomain in PARP16 [37] and perhaps other family members [38] suggest similar mechanisms of ART domain regulation in these enzymes. Finally, all family members except PARP16 possess both additional intramolecular domains and external accessory factors [4,39], with potential contribution to the regulation of catalytic activity; however, such a connection has not been demonstrated.

Here we show that the ART domain of PARP15 forms homodimers in solution (Fig. 1 – 3). Furthermore, we show that PARP15 MARylation activity is dependent on homodimerization (Fig. 4,5). This mechanism of activation has not been described for any PARP family member. Our biophysical analysis of several dimerization incompetent mutant constructs shows that these mutants are stable in solution; they are capable of ligand binding in their active sites; and they are apparently properly folded. Therefore, we believe that the gain of activity upon dimer formation is due to a physiological regulatory mechanism and not merely a stabilization of the structure. We show by complementary methods that the PARP15 ART domain dimerizes at mid-nanomolar concentrations, with an approximate K_d_ of 260 nM. A binding strength in this range is quite typical for physiological protein-protein interactions [40,41].

A dimer dissociation constant in the mid-nanomolar range (Fig. 2) supports the physiological relevance of the PARP15 dimer interaction: At sufficient local concentration the domain might activate itself while maintaining the potential to be de-activated by interaction with other binding partners, or relocalization to an environment with lower concentrations. PARP15 - as TNKS1/PARP5a, PARP12, PARP13 and PARP14 - localizes to stress granules (SGs) [19], protein- and RNA-rich condensates which form in the cytoplasm when translation is halted [20]. PARP15 may dimerize upon its accumulation at SGs. As the dimeric form of PARP15 would contain four ADP-ribose binding domains its contribution to the multivalent interactions network that constitutes the liquid-like compartments would be significantly increased.

Our experimental evidence from HDX-MS and mutagenesis suggests that the solution dimer interface is consistent with the interface observed by X-ray crystallography. Our analysis of PARP15 crystal structures using the PISA server [29] gave a mean buried surface area for the crystallographic dimer interface that is in the range of similar protein interactions, including homodimers [40-43]. It is apparent that small interfaces can generate the binding free energy required for dimer formation. The PISA server calculated a free energy change of −9.9±0.6 kcal/mol upon complex formation for the PARP15 ART domains, as compared to −14.2 kcal/mol for the 14-3-3β homodimer.

In summary, this study revealed that the catalytically functional unit of PARP15 is a homodimer facilitated by its ART domain. This exposes dimerization as a novel regulatory mechanism of this domain. Next, we will need to investigate how the dimerization event is relayed to the catalytic site through the core of the protomer. Furthermore, we will need to understand the significance of this finding in the full domain context of the protein. This will be important information in gauging the value of dimerization inhibitors for pharmacological inhibition of PARP15. We anticipate that these new insights will contribute to bridging the gap between the biochemical and cellular aspects of PARP15 biology.

### Materials and Methods

#### Materials and chemicals

Unless stated otherwise, reagents and fine chemicals were purchased from Merck Sigma-Aldrich (Germany) and chromatographic media were obtained from Cytiva (Sweden).

#### Molecular Cloning

The expression vectors for the wild type human ART domains of PARP10 (Asn819-Val1007) [23], PARP14 (His1608-Lys1801) as well as wt and H599Y-PARP15 (Asn481-Ala678) [26] were generated by inserting the respective cDNAs in pNIC-Bsa4 (Genbank entry EF198106) by ligation independent cloning. The PARP15 ART wt construct was used as template in site-directed mutagenesis (Q5 site-directed mutagenesis kit, New England Biolabs) to obtain the mutant constructs R576A, R576E, R576D, D665A, and D665R. The PARP15 macrodomain-2 construct (Gly287-Asn470) in pNIC-Bsa4 was described before [15]. A synthetic gene of PARP15 macrodomain-2+ART (“m2-ART”; Gly287-Ala678), codon optimized for expression in *E*.*coli*, was obtained from GeneArt (Thermo Fisher Scientific) in pET151/TOPO containing an N-terminal 6xHis tag and V5 epitope. Note that sequence numbering in our previous publications [15,26] followed an earlier gene annotation.

#### Protein expression and purification

Expression plasmids were transformed into BL21(DE3) cells (C2527H; New England Biolabs). The cells were grown in terrific broth medium supplemented with 100 µg/ml ampicillin or 50 µg/ml kanamycin in a LEX bioreactor at 37 °C under constant aeration. After overnight induction at 18 °C with 0.4 mM isopropyl β-D-thiogalactopyranosid, cells were harvested by centrifugation at 4,600 rcf, 15 min. Harvested cells were resuspended in lysis buffer (50 mM HEPES pH 7.5, 300 mM NaCl, 10% glycerol, 0.5 mM TCEP) supplemented with protease inhibitor cocktail (Roche) and benzonase Nuclease (Merck Millipore), and cells were lysed by sonication. The lysate was clarified by centrifugation at 22,000 rcf for 25 min at 4 °C. The clarified lysate was filtered through 0.4 µm syringe filters before loading onto a HiTrap TALON crude column. Bound proteins were eluted with an imidazole gradient up to 400 mM and peak fractions were pooled and injected into a HiLoad 16/600 Superdex 75 pg column, equilibrated in 50 mM HEPES pH 7.5, 300 mM NaCl, 10% glycerol, 0.5 mM TCEP. The PARP15 m2-ART construct was instead loaded onto a HiLoad 16/600 Superdex 200 pg column, equilibrated in 50 mM NaPO_4_ pH 7.3, 0.5 mM TCEP, 5% glycerol. The pooled m2-ART elution fractions were subjected to cation exchange chromatography on a HiTrap SP XL column and PARP15 was eluted with a shallow gradient to 1M NaCl. The purity of the proteins was evaluated by SDS-PAGE. Purified proteins were aliquoted and stored at −80 °C.

#### SEC-RALS/LALS (OMNISEC)

OMNISEC runs were performed at the Lund Protein Production Platform (LP3), Sweden. 120 µl protein solution (0.75 mg/ml PARP15 constructs, 0.35 mg/ml PARP14 ART, 0.65 mg/ml PARP10 ART) was injected in duplicates into the OMNISEC system (Malvern Panalytical) equilibrated in 50 mM HEPES pH 7.5, 300 mM NaCl and 1 mM TCEP and operated at a flow rate of 0.5 ml/min. The OMISEC RESOLVE module was equipped with a Superdex 200 Increase 10/300 GL column (Cytiva). Right angle and low angle light scattering (RALS 90° angle and LALS 7° angle), differential refractive index, viscosity, and UV/VIS signal were measured by the OMNISEC REVEAL module. The integrated software (OMNISEC v11.36) was used for data collection and analysis. The refractive index and inferred molecular weight of representative runs were plotted in Prism vs. 10.1.2. (GraphPad software).

#### Chemical crosslinking

Protein at 2, 10, or 20 µM was reacted with bis(sulfosuccinimidyl)suberate (BS^3^; Merck Sigma-Aldrich, S5799) at a molar ratio of 2:1 (protein to crosslinker) in reaction buffer (50 mM HEPES pH 7.5, 100 mM NaCl, 4 mM MgCl_2_, 0.2 mM TCEP). After 10 min incubation at room temperature, the reaction was quenched with 0.4 mM Tris pH 8 and addition of Leammli buffer. The proteins were resolved on 4-12% Bis-Tris gels (ThermoFisher Scientific) and stained with Coomassie [44].

#### Mass spectrometry to analyze chemically crosslinked samples

##### In-solution digestion

Cross-linked PARP15 m2-ART construct was digested in solution: The pH of the samples was adjusted to 7.8 by addition of ammonium bicarbonate (ABC) to a final concentration of 100 mM. The proteins were reduced by addition of 5 mM DL-dithiothreitol (DTT) and incubated in 37 °C for 30 min, followed by alkylation using iodoacetamide (IAA) at 12 mM and incubation in the dark for 20 min. Sequencing-grade modified trypsin (Promega, Madison, WI, USA) was added to a concentration of 1:100 (trypsin:protein) before the samples were incubated at 37 °C for four hours. A second aliquot of trypsin was added giving a final concentration of 1:50 (trypsin:protein) and the samples were incubated over night at 37 °C. The next day, formic acid (FA) was added to a final concentration of 0.5 %.

##### In-gel digestion

Gel bands were excised from SDS-PAGE gels, cut into 1×1 mm pieces and transferred to 1.5 ml tubes. The gel pieces were washed and incubated for 30 min three times with 500 µl 50% acetonitrile (ACN), 50 mM ABC. The gel was dehydrated using 100% ACN before the proteins were reduced with 25 µl 10 mM DTT in 50 mM ABC for 30 min at 37 °C. DTT was removed and the gel was dehydrated again using 100% ACN before the proteins were alkylated with 25 µl 55 mM iodacetamide in 50 mM ABC for 30 min in the dark at room temperature. The gel was dehydrated one last time with 100 % ACN before the proteins were digested by adding 25 µl 12 ng/µl trypsin (sequence grade modified porcine trypsin, Promega, Fitchburg, WI, USA) in 50 mM ABC and incubated on ice for 4 hours before 20 µl 50 mM ABC were added and the protein were incubated overnight at 37 °C. The following day, formic acid was added to a final concentration of 0.5 %, to get a pH of 2-3, before the peptide solutions were extracted and transferred into new tubes.

##### LC-MSMS

The peptides were cleaned up using C18 reversed-phase micro columns with a 2% ACN, 0.1% FA equilibration buffer and an 80% ACN, 0.1% FA elution buffer. The collected samples were dried in a fume hood and resuspended in 15 µl 2% ACN, 0.1% FA. The samples were then injected to an ultra-high pressure nanoflow chromatography system (nanoElute, Bruker Daltonics). The peptides were loaded onto an Acclaim PepMap C18 (5 mm, 300 μm id, 5 μm particle diameter, 100 Å pore size) trap column (Thermo Fisher Scientific) and separated on a Bruker Pepsep Ten C18 (75 µm × 10 cm, 1.9 µm particle size) analytical column (Bruker Daltonics). Mobile phase A (2% ACN, 0.1% FA) was used with the mobile phase B (0.1% FA in ACN) for 45 min to create a gradient (from 2 to 17% B over 20 min, from 17 to 34% B over 10 min, from 34 to 95% B over 3 min, at 95% B over 12 min) at a flow rate of 400 nl/min and a column oven temperature of 50 °C. The peptides were analysed on a quadrupole time-of-flight mass spectrometer (timsTOF Pro, Bruker Daltonics), via a nano electrospray ion source (Captive Spray Source, Bruker Daltonics) in positive mode, controlled by the OtofControl 5.1 software (Bruker Daltonics). The temperature of the ion transfer capillary was 180°. A DDA method was used to select precursor ions for fragmentation with one TIMS-MS scan and 10 PASEF MS/MS scans. The TIMS-MS scan was acquired between 0.60–1.6 V s/cm^2^ and 100–1700 m/z with a ramp time of 100 ms. The 10 PASEF scans contained maximum of 10 MS/MS scans per PASEF scan with a collision energy of 10 eV. Precursors with maximum 5 charges with intensity threshold to 5000 a. u. and a dynamic exclusion of 0.4 s were used.

##### Data analysis

Raw data were processed using Mascot Distiller (version 2.8) and searched against an in-house database containing the PARP15 protein sequence using the settings; precursor ion tolerance 8 ppm, MS/MS fragment mass tolerance 0.015 Da, trypsin as protease, 1 missed cleavages site, methionine oxidation as variable modification and carbamidomethylation as fixed modification. The obtained peak lists were analyzed in xiSEARCH [45]. As both in-solution and in-gel digest data obtained similar results, three peak lists of each method were pooled and analyzed together to improve stringency of the results. The following parameters were applied: crosslinker = BS^3^(small scale); enzyme = trypsin; misscleavages = 3; fixed modifications = carbamidomethylation (C); variable modifications = oxidation (M); variable (linear peptides) = BS^3^amidated, BS^3^Hydrolized; ions = b-ion, y-ion. The output was visualized in xiVIEW [46], where low-scoring peptides were filtered out.

#### Mass photometry

The concentration-dependent dimerization of PARP15 constructs was followed on a TwoMP mass photometer (REFEYN, Oxford, UK). Microscopy glass slides (24 × 50 mm) were alternately cleaned with ultrapure water and isopropanol and finally dried under a clean nitrogen stream. Silicone gasket wells (Grace Biolabs, Merck Life Science AB, Solna, Sweden) were placed on the clean glass slide and the focus was adjusted to a well containing 15 µl sample buffer (20 mM HEPES pH 7.5, 50 mM NaCl, 4 mM MgCl2, 0.5 mM TCEP). The proteins were added to the sample buffer at varying concentrations (see Fig. 1H) and landing events were recorded over 60 seconds in a frame size of 900 × 354 pixels and at a frame rate of 496 Hz. The DiscoverMP software provided by the manufacturer was used for data analysis. To translate recorded mass photometry contrast into molecular weight, a native PAGE marker was used for calibration (Invitrogen, Thermo Fisher Scientific, Gothenburg, Sweden).

#### Fluorescence polarization

PARP15 ART was diluted to a concentration of 2 mg/ml in 0.1 M sodium carbonate-bicarbonate buffer pH 9.0 in a light-proof safe-lock tube. Fluorescein isothiocyanate (Merck Sigma-Aldrich, F4274) was dissolved in 0.1 M sodium carbonate-bicarbonate buffer pH 9.0 at a concentration of 1 mg/ml. A 10-fold molar access of FITC was then added dropwise to the vial containing the protein solution. The reaction was allowed to proceed in the dark under constant shaking for 2 h at room temperature. Unreacted FITC was removed with a PD SpinTrap G-25 column (Cytiva, #28918004) equilibrated in 1x phosphate buffered saline. The protein concentration and degree of labelling were determined by spectrophotometry.

Fluorescence polarization assays were performed in 96-well half-area non-binding plates (Corning, #3993) in 50 mM HEPES pH 7.5, 100 mM NaCl, 4 mM MgCl2, 0.2 mM TCEP, 0.01% NP-40. A three-fold dilution series of unlabelled PARP15 ART was prepared before adding 1 nM or 10 nM FITC-labelled PARP15 to each well, resulting in a final sample volume of 80 µl per well. Biological triplicates of each reaction were incubated for 15 min at room temperature, gently shaking (250 rpm) to allow for the system to equilibrate. Fluorescence polarization was recorded on a CLARIOStar multimode reader (BMG Labtech) in top-optic, end-point mode with a 482-16 nm excitation filter, a 530-40 emission filter, and an LP 504 nm dichroic filter. To obtain the fraction of FITC-PARP15 in a dimeric state at a given protein concentration, we used the formula

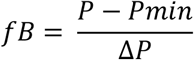

where fB is the fraction bound, P is the measured polarization, Pmin is the minimum polarization, and ΔP is (Pmax-Pmin) the total change in polarization. The fractions obtained were plotted against protein concentration and dissociation constants were determined by fitting the data to a one-site specific binding equation using in Prism vs. 10.1.2. (GraphPad software).

#### MacroGreen assay

ADP-ribosylation activity of PARPs was determined using a green fluorescent protein fusion of a modified Af1521 macrodomain [33]. For auto-ADP-ribosylation assays, a 2:1 dilution series (two parts protein, one part buffer) in assay buffer (50 mM HEPES pH 7.5, 100 mM NaCl, 4 mM MgCl2, 0.2 mM TCEP) starting from 1 µM PARP protein was prepared in a 96-well flat-bottom plate. The plate was incubated for 20 min at room temperature to allow for a monomer-dimer equilibrium to be reached. Reactions were started by addition of 120 µM NAD+. ADP-ribosylation reactions were performed at room temperature for 30 min under constant shaking at 250 rpm. The content of each well was distributed into three individual wells on a MaxiSorp™ plate (NUNC). The proteins were allowed to bind to the plate for an additional 15 min (at room temperature, 250 rpm). When PARP15 macrodomain-2 was used as a substrate, 250 nM PARP15 ART wt, D665A, or R576A, was mixed with titrations of PARP15 ART H559Y in 50 mM HEPES pH 7.5, 100 mM NaCl, 4 mM MgCl2, 0.2 mM TCEP (2-fold dilution series of H559Y starting from 2 µM). PARP15 ART H559Y was omitted in the control reaction and another set of reactions containing only H559Y were set up to show that the mutant is indeed inactive. An equilibrium was allowed to establish for 1 h at room temperature and 250 rpm shaking. Meanwhile, the wells of a MaxiSorp™ plate (NUNC) were coated with PARP15 macrodomain-2 for 30 min at room temperature, 250 rpm, and subsequently blocked with 1% BSA in reaction buffer for an additional 15 min. The wells were rinsed twice with 150 µl reaction buffer before 50 µl of the PARP15 ART-containing samples were added to the MaxiSorp™ plate and the ADP-ribosylation reactions were started by the addition of 100 µM NAD+. The reaction was allowed to proceed for 40 min at room temperature, 250 rpm. For detection, 1 µM MacroGreen was added to the wells. Fluorescence intensity was measured in a CLARIOStar multimode reader (BMG Labtech) using a 470-15 nm extinction filter and a 515-20 nm emission filter. Raw data were plotted in Prism vs. 10.1.2. (GraphPad software).

#### Hydrogen-Deuterium Exchange Mass Spectrometry

##### Experiment

pH measurements were made using a SevenCompact pH-meter equipped with an InLab Micro electrode (Mettler-Toledo), a 4-point calibration (pH 2,4,7,10) was made prior to all measurements. The HDX-MS analysis was made using automated sample preparation on a LEAP H/D-X PAL™ platform interfaced to an LC-MS system, comprising an Ultimate 3000 micro-LC coupled to an Orbitrap Q Exactive Plus MS. HDX was performed on 2 mg/ml PARP15 in 50 mM TBS, pH 7.4, for each timepoint 3 µl HDX samples were diluted with 27 µl labelling buffer (50mM TBS) prepared in D2O, pH_(read)_ 7.0. The HDX labelling was carried out for t = 0, 30, 300, 3000 and 9000s at 20 °C. The labelling reaction was quenched by dilution of 30 µl labelled sample with 30 µl of 1% TFA, 0.4 M TCEP, 4 M urea, pH 2.5 at 1°C, 60 µl of the quenched sample was directly injected and subjected to online pepsin digestion at 4 °C (in-house immobilized pepsin column, 2.1 × 30 mm). Online digestion and trapping was performed for 4 minutes using a flow of 50 µl/min 0.1 % formic acid, pH 2.5. The peptides generated by pepsin digestion were subjected to on-line SPE on a PepMap300 C18 trap column (1 mm × 15mm) and washed with 0.1% FA for 60s. Thereafter, the trap column was switched in-line with a reversed-phase analytical column, Hypersil GOLD, particle size 1.9 µm, 1 × 50 mm, and separation was performed at 1°C using a gradient of 5-50 % B over 8 minutes and then from 50 to 90% B for 5 minutes, the mobile phases were 0.1 % formic acid (A) and 95 % acetonitrile/0.1 % formic acid (B). Following the separation, the trap and column were equilibrated at 5% organic content, until the next injection. The needle port and sample loop were cleaned three times after each injection with mobile phase 5% MeOH/0.1% FA, followed by 90% MeOH/0.1% FA and a final wash of 5% MeOH/0.1% FA. After each sample and blank injection, the pepsin column was washed by injecting 90 ul of pepsin wash solution 1% FA /4 M urea /5% MeOH. To minimize carry-over, a full blank was run between sample injections. Separated peptides were analysed on a Q Exactive Plus MS, equipped with a HESI source operated at a capillary temperature of 250 °C with sheath gas 12, Aux gas 2 and sweep gas 1 (au). For HDX analysis MS full scan spectra were acquired at 70K resolution, AGC 3e6, Max IT 200ms and scan range 300-2000. For identification of generated peptides separate undeuterated samples were analysed using data dependent MS/MS with HCD fragmentation.

##### Data analysis

PEAKS Studio × Bioinformatics Solutions Inc. (BSI, Waterloo, Canada) was used for peptide identification after pepsin digestion of undeuterated samples. The search was done on a FASTA file with only the RDB sequence, search criteria was a mass error tolerance of 15 ppm and a fragment mass error tolerance of 0.05 Da, allowing for fully unspecific cleavage by pepsin. Peptides identified by PEAKS with a peptide score value of log P > 25 and no modifications were used to generate a peptide list containing peptide sequence, charge state and retention time for the HDX analysis. HDX data analysis and visualization was performed using HDExaminer, versio 3.1.1 (Sierra Analytics Inc., Modesto, US). The analysis was made on the best charge state for each peptide, allowed only for EX2 and the two first residues of a peptide was assumed unable to hold deuteration. Due to the comparative nature of the measurements, the deuterium incorporation levels for the peptic peptides were derived from the observed relative mass difference between the deuterated and non-deuterated peptides without back-exchange correction using a fully deuterated sample [47]. As a full deuteration experiment was not done, full deuteration was set to 75% of maximum theoretical uptake. The deuteration data presented are the average of all high and medium confidence results. The allowed retention time window was ± 0.5 minute. The spectra for all time points were manually inspected; low scoring peptides, obvious outliers and any peptides where retention time correction could not be done consistently were removed. As bottom-up labelling HDX-MS is limited in structural resolution by the degree of overlap of the peptides generated by pepsin digestion, the peptide map overlap is shown for each respective state in Supplementary Figure S2.

#### Bioinformatic analysis and structure visualization

ColabFold v1.5.5 [31] was used for predictions of the PARP15 m2-ART structure and PARP15 ART homodimer. Crystal structures of the PARP15 ART domain deposited in the Protein Data Bank (PDB) were analyzed using the PISA server [29]. PyMOL was used for structural alignments and general representation of experimental structures and computational models.

#### Analytical size exclusion chromatography

A Superdex 200 Increase 10/300 GL column was equilibrated in running buffer (50 mM HEPES pH 7.5, 300 mM NaCl, 10% glycerol, 0.5 mM TCEP) and calibrated using the following standards: Ovalbumin (44 kDa, 2.5 mg/ml), conalbumin (75 kDa, 2.5 mg/ml), aldolase (158 k Da, 2.5 mg/ml), ferritin (440 k Da, 2.5 mg/ml), thyroglobulin (669 kDa, 2.5 mg/ml), RNase (13.7 kDa, 3.2 mg/ml), Blue Dextran (2.5 mg/ml). The column was operated at a flow rate of 0.7 ml/min on an ÄKTA pure FPLC. Subsequently, 100 µl of each PARP15 ART construct at 0.7 mg/ml were injected into the column, using the same run parameters as before. Chromatograms were exported from the Uniprot software, analyzed in Microsoft Excel, and re-plotted using Prism vs. 10.1.2. (GraphPad software).

#### Biotin-NAD western-blot-based assay

10 µM of PARP15 ART constructs or, 5 µM in case of mutant combinations, were incubated in assay buffer (20 mM HEPES pH 7.5, 50 mM NaCl, 4 mM MgCl_2_, 0.2 mM TCEP) for 30 min at 37°C, 250 rpm, prior to the addition of NAD+. 50 µM NAD+ containing 10% biotin-NAD+ were added to start the ADP-ribosylation reaction, followed by incubation for another 30 min. The reactions were stopped by the addition of Leammli buffer and heating for 1 min at 95°C. Samples were then loaded onto NuPage 4-12% Bis-Tris gels and resolved by electrophoresis. Proteins were transferred from the gel onto a PVDF membrane in a wet-transfer set-up, using a transfer buffer consisting of 3 g/L Trisma-Base, 14.4 g/L glycine, and 10% methanol. The transfer was performed for 60 min at 25 V. Ponceau S staining of the membrane was used to assess transfer efficiency and served as a loading control. The membrane was blocked with 1% BSA in TBS-T buffer for 20 min and then incubated in 0.5 µg/ml Streptavidin-HRP (Pierce. PIER21126) in 1% BSA in TBS-T buffer for 35 min. For detection, we used Clarity Western ECL substrate (BIORAD) following the manufacturer’s instructions. Chemiluminescent and colorimetric images were recorded on a ChemiDoc Imaging System (BIORAD).

#### Differential Scanning Fluorimetry

The PARP15 ART domains were diluted to 0.2 mg/ml in 50 mM HEPPES pH 7.5, 100 mM NaCl, 4 mM MgCl2, 0.2 mM TCEP. 1 mM 3-aminobenzamide was added and the samples were incubated at room temperature for at least 45 min. Three Prometheus NT.48 standard capillaries (Nanotemper) were loaded per sample and the intrinsic fluorescence at 330 nm and 350 nm was recorded over a temperature gradient of 25 – 95 °C (+1 °C/min) in a Prometheus Pantha (NanoTemper). Raw data, first derivatives and melting temperatures were exported from the PR.ThermControl software and plotted in Prism vs. 10.1.2. (GraphPad software). The melting temperatures of the mutants were compared to wt PARP15 using a one-way ANOVA and the pairwise significance was determined using a Dunnett’s multiple comparisons test.

## Acknowledgements

We thank Céleste Sele at the Lund University Protein Production Platform (LP3) Sweden for OMNISEC analyses and assistance with nanoDSF, and Anders Hofer and Saber Anoosheh (Umeå University, Sweden) for their introduction to mass photometry.

## Author Contributions

C.E.: Conceptualization, experiment, data analysis, visualization, manuscript writing. A.G.G.S.: Experiment, data analysis. S.E.: Experiment, data analysis. K.B.: Experiment, data analysis. H.S.: Conceptualization, manuscript writing, funding acquisition.

