## Supplemental Information Document for "Regulation of ADP-ribosyltransferase activity by ART domain dimerization in PARP15"

### Supplementary Data

**Supplementary Table S1.** Details on the SEC-RALS/LALS analysis of indicated PARP constructs.

| | PARP15 ART | PARP14 ART | PARP10 ART | | PARP15 m2 | PARP15 m2-ART |
|---|---|---|---|---|---|---|
| | | | Peak 1 | Peak 2 | | |
| ***RV (ml)** | 16.02 | 17.34 | 15.79 | 16.61 | 16.66 | 14.47 |
| **Mw (g/mol)** | 47 678 | 25 903 | 54 347 | 31 636 | 23 071 | 71 361 |
| **Mp (g/mol)** | 47 253 | 25 695 | 52 942 | 30 210 | 22 365 | 71 604 |
| **Calculated Mw (g/mol)** | 25 402 | 24 966 | 24 034 | 24 034 | 22 382 | 47 316 |
| **Mw/Mn** | 1.001 | 1.001 | 1.003 | 1.005 | 1.001 | 1.003 |
| **IVw (dL/g)** | 0.0314 | 0.0219 | - | 0.0214 | 0.0419 | 0.0462 |
| **Rhw (nm)** | 2.89 | - | - | 2.17 | 2.48 | 3.73 |
| **Frac. Of sample (%)** | 100 | 100 | 9 | 91 | 100 | 100 |
| **Measured conc. (mg/ml)** | 0.65 | 0.37 | 0.49 | 0.49 | 0.74 | 0.55 |
| **Recovery (%)** | 87.0 | 106 | 74.7 | 74.7 | 98.4 | 73.3 |

*RV: retention volume measured at peak maximum
Mw: weight average molecular weight from the molecular weight distribution
Mp: molecular weight at the peak apex
Calculated Mw: calculated using http://web.expasy.org/protparam/ based on the amino acid sequence of the expression construct
Mw/Mn: dispersity
IVw: weight average intrinsic viscosity
Rhw: weight average hydrodynamic radius
Frac. of sample: fraction of the sample in this peak expressed as a percent of the total area of all the analyzed peaks
Measured conc.: measured concentration of the peak
Recovery: recovery of sample based on RI peak area and input dn/dc compared to the input sample concentration
Definitions from the OMNISEC System User Guide MAN0550-06-EN

**Supplementary Table S2**. Selected details of an interface analysis of a PARP15 ART domain crystal dimer using the PISA server (1)

| PDB entry 6ry4 (2) | Chain A | Chain B |
|---|---|---|
| **No. Interface residues** | 27 (13.7%) | 27 (13.7%) |
| **Solvent accessible area (Å2)** | 1051.3 (9.9%) | 1061.4 (9.9%) |
| **Gain of solvation energy on complex formation (kcal/mol)** | -4.8 | -5.8 |

**Interface details**

| # | Protomer 1 | Dist. [Å] | Protomer 2 |
|---|---|---|---|
| | **Hydrogen Bonds** | | |
| **1** | B: GLN 543 [NE2] | 3.11 | A: PHE 532 [O] |
| **2** | B: GLN 543 [NE2] | 3.06 | A: GLN 535 [OE1] |
| **3** | B: GLN 543 [NE2] | 3.60 | A: SER 536 [OG] |
| **4** | B: ARG 576 [NH2] | 2.94 | A: ASN 575 [OD1] |
| **5** | B: ARG 576 [NE] | 2.94 | A: SER 577 [OG] |
| **6** | B: HIS 572 [NE2] | 2.73 | A: THR 644 [O] |
| **7** | B: ARG 576 [NH2] | 2.97 | A: ASP 665 [OD1] |
| **8** | B: ASP 665 [N ] | 2.76 | A: ASP 665 [OD2] |
| **9** | B: ARG 576 [NH1] | 2.92 | A: ASP 665 [OD2] |
| **10** | B: PHE 532 [O] | 3.04 | A: GLN 543 [NE2] |
| **11** | B: GLN 535 [OE1] | 2.96 | A: GLN 543 [NE2] |
| **12** | B: SER 536 [OG] | 3.66 | A: GLN 543 [NE2] |
| **13** | B: ASN 575 [OD1] | 2.86 | A: ARG 576 [NH2] |
| **14** | B: SER 577 [OG] | 2.73 | A: ARG 576 [NE] |
| **15** | B: THR 644 [O] | 2.58 | A: HIS 572 [NE2] |
| **16** | B: ASP 665 [OD1] | 2.96 | A: ARG 576 [NH2] |
| **17** | B: ASP 665 [OD2] | 2.74 | A: ASP 665 [N] |
| **18** | B: ASP 665 [OD2] | 3.11 | A: ARG 576 [NH1] |
| | **Salt Bridges** | | |
| **1** | B: ARG 576 [NH1] | 3.75 | A: ASP 665 [OD1] |
| **2** | B: ARG 576 [NH2] | 2.97 | A: ASP 665 [OD1] |
| **3** | B: ARG 576 [NH1] | 2.92 | A: ASP 665 [OD2] |
| **4** | B: ARG 576 [NH2] | 3.56 | A: ASP 665 [OD2] |
| **5** | B: ASP 665 [OD1] | 3.74 | A: ARG 576 [NH1] |
| **6** | B: ASP 665 [OD1] | 2.96 | A: ARG 576 [NH2] |
| **7** | B: ASP 665 [OD2] | 3.11 | A: ARG 576 [NH1] |
| **8** | B: ASP 665 [OD2] | 3.67 | A: ARG 576 [NH2] |

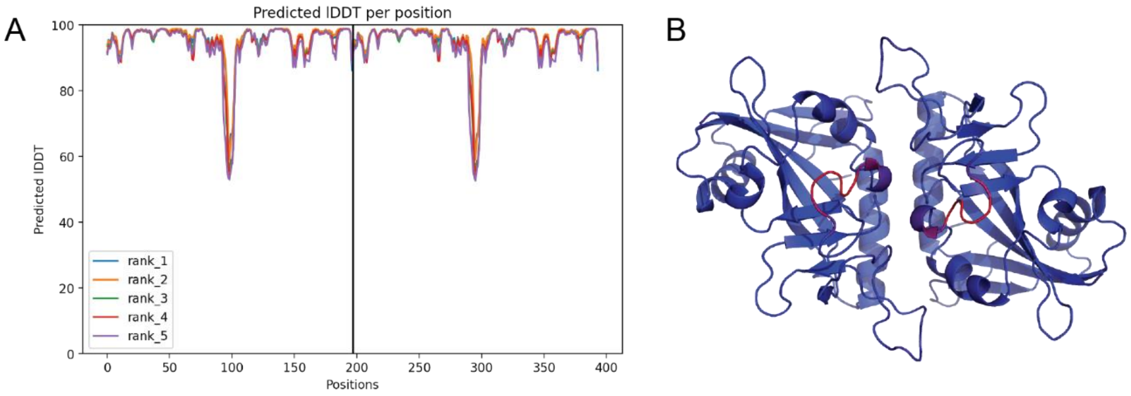

**Supplementary Figure S1. Quality of the Alphafold2 multimer model of the PARP15 ART domain dimer.** (**A**) Per-residue pIDDT score (3). The two low-scoring regions represent a loop that is often not resolved in published crystal structures. (**B**) pIDDT score in (A) mapped onto the rank 1 model (blue – high confidence; red – low confidence).

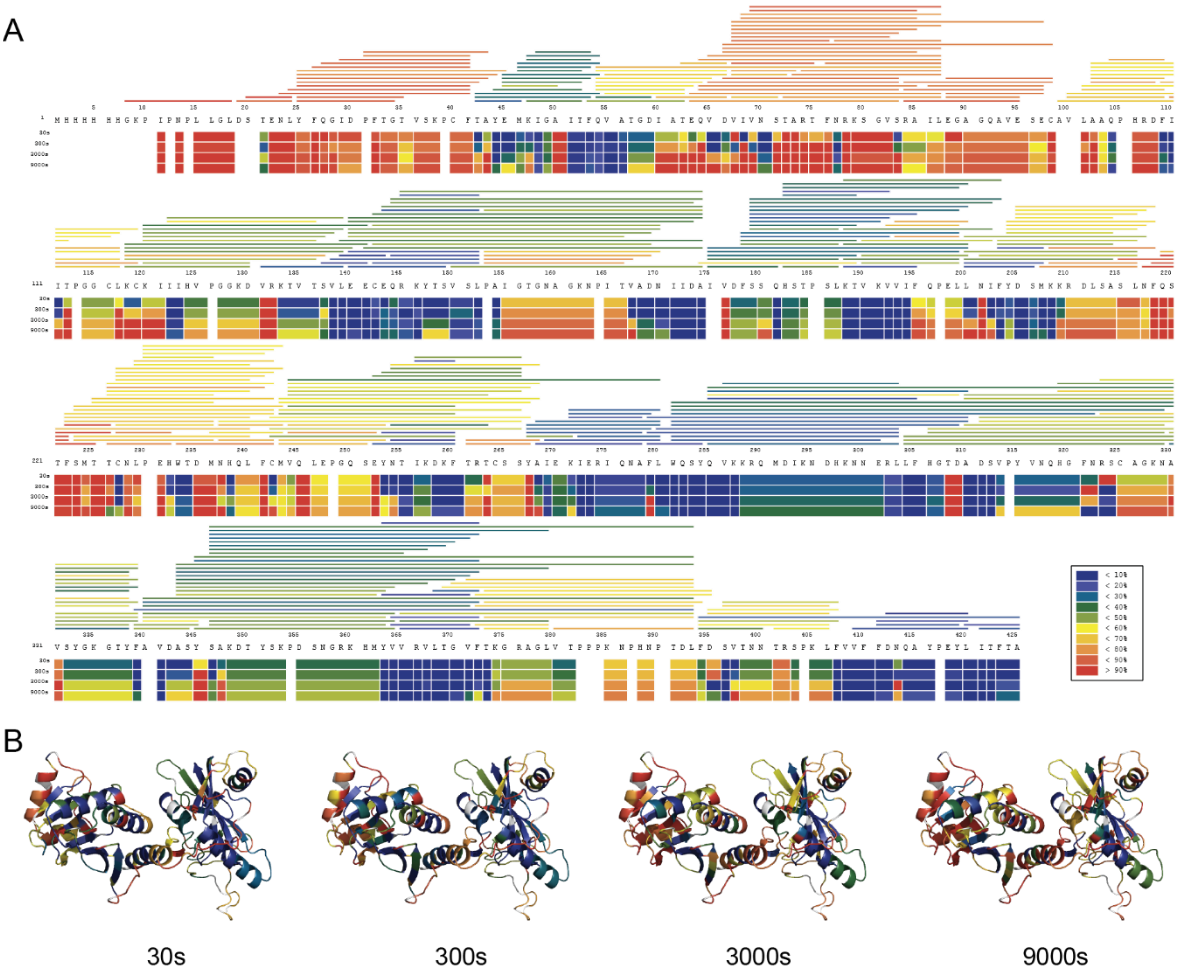

**Supplementary Figure S2. Per-peptide deuterium exchange dynamics of PARP15 m2-ART based on HDX-MS data.** (**A**) Identified peptides mapped onto the construct sequence (N-terminal His-tag, followed by TEV cleavage site, macrodomain-2, and ART domain), color coded according to percent deuterium exchange. The boxes below the sequence represent the per residue deuterium exchange based on the peptide coverage at the respective site and time point. (**B**) Data from panel A superimposed on the Alphafold2 model of the m2-ART construct (3).

**Supplementary Table S3. BS3 crosslinked peptides and their masses identified.** See main text for Materials and Methods section and further explanation.

| Pos1 | PepSeq1 | L1* | Pos2 | PepSeq2 | L2* | Score | Ch* | ExpMz | ExpMass | CalcMz | CalcMass |
|---|---|---|---|---|---|---|---|---|---|---|---|
| **581** | SCcarbamido-methyl-AGKNAVSYGK | 5 | 588 | NAVSYGKG TYFAVDASY SAK | 7 | 10.033908 | 4 | 870.17429 | 3476.668054 | 870.1712315 | 3476.65582 |
| **581** | SCcarbamido-methyl-AGKNAVSYGK | 5 | 627 | VLTGVFTKGR | 8 | 10.23641 | 4 | 614.82904 | 2455.287054 | 614.8296047 | 2455.289313 |
| **581** | SCcarbamido-methyl-AGKNAVSYGK | 5 | 606 | DTYSKPDSNGR | 5 | 11.601239 | 4 | 655.30822 | 2617.203774 | 655.3092332 | 2617.207827 |
| **581** | SCcarbamido-methyl-AGKNAVSYGK | 5 | 581 | SCcarbamido-methyl-AGKNAVSYGK | 5 | 11.407647 | 4 | 655.81485 | 2619.230294 | 655.8178032 | 2619.242107 |
| **581** | SCcarbamido-methyl-AGKNAVSYGK | 5 | 627 | VLTGVFTKGR | 8 | 10.697531 | 3 | 819.43741 | 2455.290401 | 819.4370475 | 2455.289313 |
| **581** | SCcarbamido-methyl-AGKNAVSYGK | 5 | 606 | DTYSKPDSNGR | 5 | 11.759573 | 3 | 873.41098 | 2617.211111 | 873.4098855 | 2617.207827 |

*L1, Link position, peptide 1; L2, Link position, peptide 2; Ch, Charge

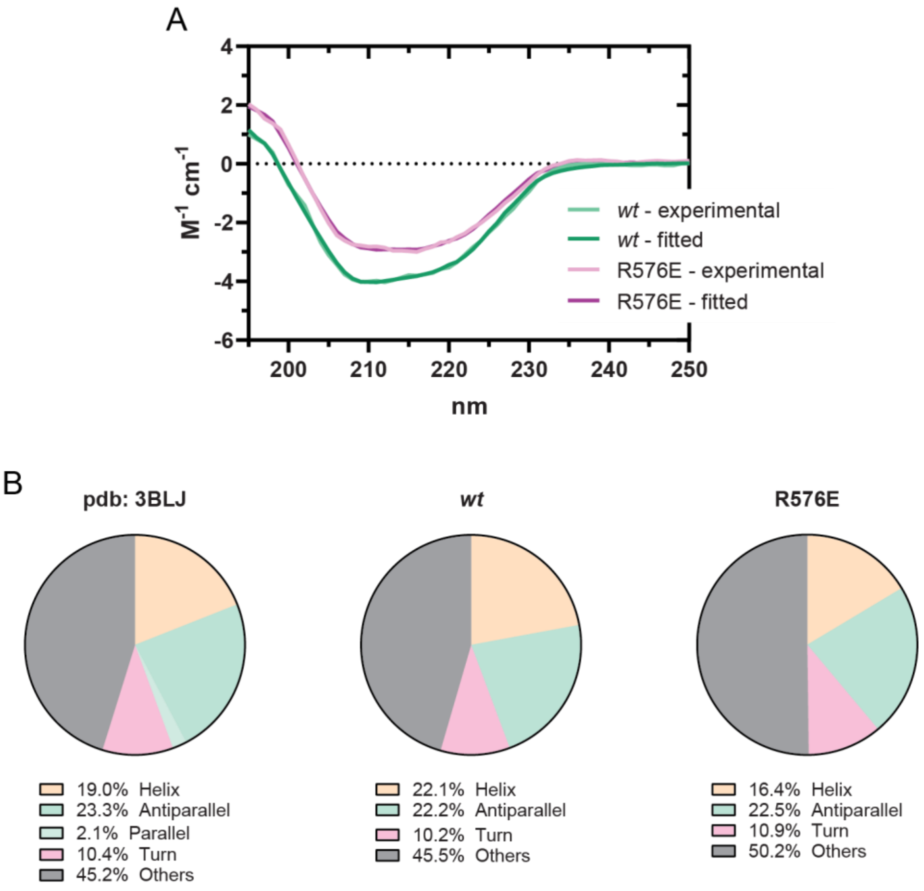

**Supplementary Figure S3. Circular dichroism spectroscopy analysis of the wild type ART domain and the interface mutant R576E.** (**A**) CD spectra for wild type and R576E ART domain. (**B**) Schematic representation of the secondary structure content of a PARP15 ART domain structural model based on X-ray crystallography (PDB entry 3BLJ) (4) as well as the data from panel A for wild type and R576E ART domain. Data were extracted and quantified using BeStSel (5).

A

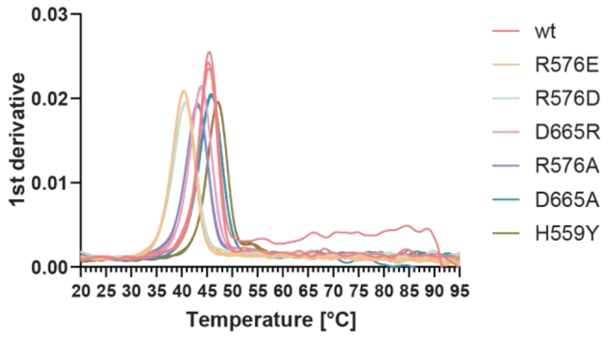

B

| variant | Tm (apo) | | Tm (3ab) | |
|---|---|---|---|---|
| | mean | SD | mean | SD |
| wt | 45.3°C | 0.09°C | 48.3°C | 0.03°C |
| R576E | 40.4°C | 0.04°C | 43.6°C | 0.03°C |
| R576D | 40.7°C | 0.06°C | 43.6°C | 0.05°C |
| D665R | 43.8°C | 0.02°C | 46.8°C | 0.08°C |
| R576A | 43.1°C | 0.02°C | 46.0°C | 0.06°C |
| D665A | 45.8°C | 0.03°C | 48.4°C | 0.02°C |
| H559Y | 47.1°C | 0.03°C | 47.5°C | 0.04°C |

C

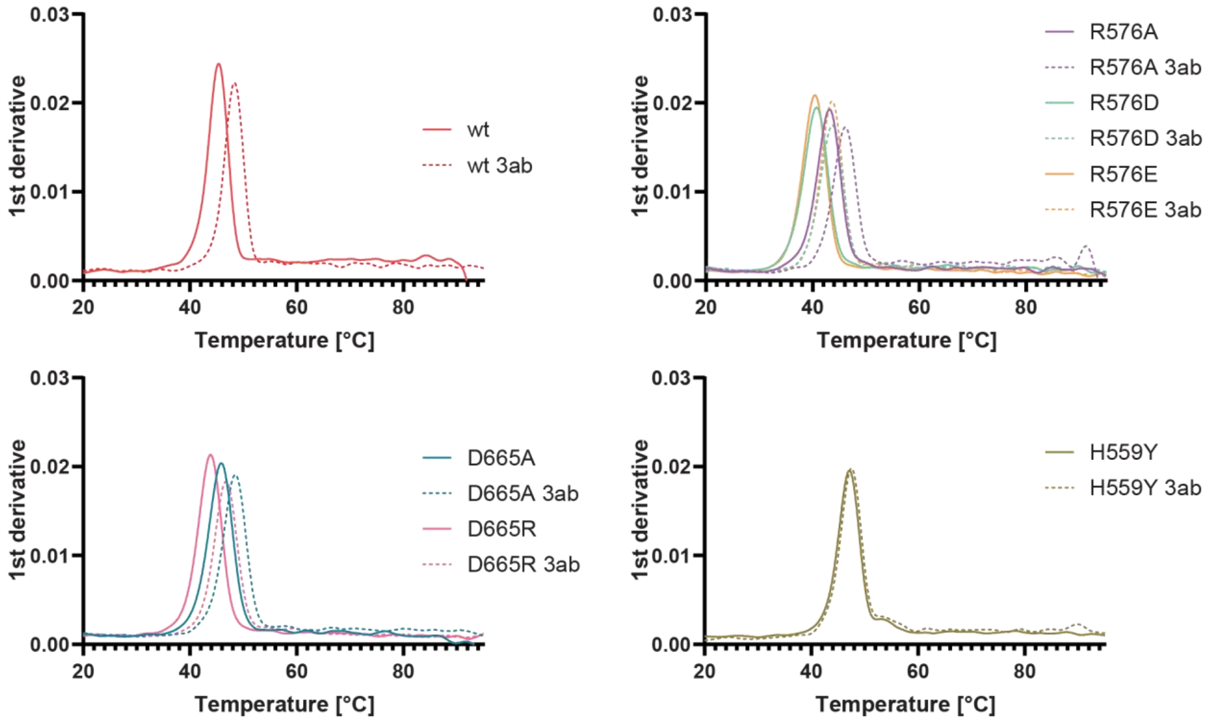

**Supplementary Figure S4. Thermal stability analysis of ART domain dimer interface mutants by nanoDSF.** (A) First derivatives ofeach variant, including the wt ART domain and the catalytically inactive H559Y mutant. Most interface mutants were less stable than the wild type ART domain, but melted in symmetrical peaks. Three replicates are shown. (B) Mean melting temperatures and standard deviations (in degrees Celsius) derived from panel (A) and (C). (C) shows the averaged first derivatives of the ART variants, with and without the addition of 3-aminobenzamide.

### Supplementary Materials and Methods

#### Circular Dichroism Spectroscopy

PARP15 ART *wt* and R576E were first desalted into assay buffer (20 mM Sodium-Phosphate buffer pH 7.5, 0.2 mM TCEP) using PD SpinTrap G-25 columns. The proteins were diluted to 0,058 mg/ml (*wt*) and 0,5 mg/ml (R576E) and transferred to a 0,1 cm quartz cuvette (Hellma; 110-1-40). The CD signal (mdeg) was recorded on a JASCO J-815 Circular Dichroism Spectrometer over a wavelength range of 250-185 nm with a data pitch of 1 nm, a scanning speed of 20 nm/min, and a bandwidth of 1 nm. For each sample, three measurements were performed and averaged. A measurement with only assay buffer was performed to determine the baseline signal which was subtracted from the raw data during data analysis. To estimate the secondary structure content of the proteins, BeStSel (5) was used to fit the raw data in millidegrees. Based on the goodness of the fit, we truncated the data in the short wavelength range and used only the data from 195-250 nm for the fitting, improving the confidence of the secondary structure estimation. The BeStSel output (mean residue ellipticity, fit to data, and percent secondary structure elements) was re-plotted in Prism vs. 10.1.2. (GraphPad software). BeStSel was also used to determine the secondary structure content of existing crystal structures using the same algorithm.
